# gcat, its_a_scam: modular tools for single cell data processing and per-cell quality control

**DOI:** 10.64898/2026.11.03.764034

**Authors:** Rishvanth K. Prabakar, Hannah V. Meyer

## Abstract

**Motivation:** The era of designing bioinformatic pipelines using open source tools is giving way to monolithic, and often proprietary, tools designed for end-to-end analysis, especially in the context of single cell data. While these tools are convenient for going from sequencing data to an almost complete data analysis “in a few clicks”, they can strongly limit data use beyond standard analyses.

**Results:** We developed *Gene and Chromosome Analysis Toolkit* (gcat) and *It’s a scam To Sell Another Single Cell Analysis Method* (`its_a_scam`) for modular processing and quality control of single cell genomic data, freeing users from their reliance on proprietary tools. gcat is a C++ library for parsing genomic data and working with it over genomic coordinates. `its_a_scam` is built on top of gcat and provides a suite of tools for a wide range of single cell data processing and quality control. `its_a_scam` can be used in conjunction with commonly used open source tools for scRNA-seq and scATAC-seq count matrix generation, and for performing per-cell quality control directly from alignment files. These alignment-level per-cell metrics can be used for quality control of each cell or to understand celltype differences post-clustering or post-annotation. Using a peripheral blood mononuclear cell Multiome dataset, we show that the count matrices generated with the aid of `its_a_scam` are highly concordant with those generated by proprietary pipelines. The alignment metrics further demonstrate celltype differences in aligned genomic regions, metagene profiles, and genebody coverage. The modular nature of `its_a_scam` allows users to integrate it into their existing pipelines, while also enabling analysis beyond current pipelines.

**Availability:** gcat (distributed as gcatlib) and `its_a_scam` are open source and available at https://github.com/rishvanth-kp/gcatlib and https://github.com/rishvanth-kp/its_a_scam. Detailed documentation is available at https://gcatlib.readthedocs.io/ and https://its-a-scam.readthedocs.io/.

## Introduction

In the age of using convenient tools for end-to-end analysis of single cell data, why do we need *Gene and Chromosome Analysis Toolkit* (`gcat`) and *It’s a scam To Sell Another Single Cell Analysis Method* (`its_a_scam`)? Single cell analysis has revolutionized our understanding of heterogeneity in celltypes [1, 2]. Building on this, pooled analysis of data across studies has enabled the creation of cell atlases and aided in the discovery of rare celltypes often undetected in individual studies [3]. However, a key challenge in such pooled analysis is to use a uniform pipeline for generating count matrices and performing quality control. Simply using the count matrices directly from public repositories introduces an unnecessary source of technical variation, because they are often generated with different tools or different versions of the same tool [4, 5]. Despite this, an open-source modular pipeline facilitating uniform quality control (QC) and analysis across the wide array of underlying protocols has been lacking. `its_a_scam`, built on top of `gcat`, provides modular tools that integrate into existing pipelines to produce count matrices concordant with those from established tools. Unlike the often inflexible proprietary tools, `its_a_scam` enables analysis and generation of per-cell quality control metrics from alignment files that are typically not possible to generate from count matrices. Furthermore, advanced users can leverage `gcat` to develop custom analysis tools.

## Results

### gcat: C++ primitives for annotations and genomic regions

`gcat` provides C++ class interfaces for parsing FASTQ, SAM/BAM, BED, and GTF files, as well as data structures for storing information over genomic coordinates (Table 1). It is a lightweight library depending only on HTSlib [6] for accessing compressed alignment files. Our rationale for choosing C++ is the fast execution of the large number of per-cell tasks and bookkeeping, work that multiplies with the number of cells. Furthermore, a compiled standalone binary file enables easy integration into existing pipelines and workflows.

**Table 1.** `gcat` **provides C++ classes for parsing genomic data and working with genomic coordinates**.

| Class name | Brief description |
|---|---|
| *File reading and parsing* |  |
| FastqReader | Takes a file name as input and reads each entry into a FastqEntry. In paired-end mode, two FastqEntry objects are returned, one from each file. |
| FastqEntry | Stores the four FASTQ fields as strings. |
| SamReader | Takes a file name as input and reads each alignment line into a SamEntry for single-end reads. For paired-end reads, a read and its mate are returned as a pair of SamEntry objects (by keeping track of each alignment until its mate is encountered). |
| SamEntry | Stores all fields of a SAM/BAM alignment entry in the appropriate data types. Also provides helper functions for parsing the SAM flag and CIGAR string. |
| BedReader | Takes a file name as input and reads the genomic coordinates into a GenomicRegion and, option-ally, all the remaining fields into a vector. |
| GenomicRegion | Stores the chromosome, start, and end of a genomic region. |
| GtfReader | Takes a file name as input and parses each line of a generic GTF file into a GtfEntry or a Gencode GTF file into a GencodeGtfEntry. |
| GtfEntry | Stores the mandatory GTF fields in the appropriate data types and the optional tag-value pairs in a hash table. |
| GencodeGtfEntry | Stores the mandatory GTF fields and the Gencode attributes in the appropriate data types. |
| *Data structures* |  |
| GenomicStepVector | A chromosome-aware and memory-efficient data structure for storing a step function in genomic coordinate space. The stored value type is a template parameter, and accumulation is defined by that type. The stored values can be retrieved at a single position or over a range of positions. |
| FeatureVector | A container type for use with GenomicStepVector. Accumulation appends to a vector so that a region can hold multiple features. |
| Metagene | Generates normalized genomic locations over variable-length non-contiguous genomic regions for creating metagene plots. |

Python-based HTSeq introduced a common (`StepVector`) and chromosome-aware (`GenomicArray`) data structure for representing a step function over coordinates by storing only the value change positions [7, 8]. gcat provides a templated C++ implementation of `GenomicArray`, called `GenomicStepVector`, for use with arbitrary data types associated with a genomic region including integers/floating point numbers, strings, or arrays of strings. `GenomicStepVector` is used in `its_a_scam` as a queryable index for both unique and overlapping annotations, and as an accumulator for coverage. Accumulation over an interval is defined by the data type a `GenomicStepVector` is instantiated with – a numeric type adds to an existing value, and `gcat’s FeatureVector` appends to an array of features, making overlapping annotations possible. `gcat` provides a Metagene class that uses a `GenomicStepVector` to generate a contiguous normalized coordinate space derived from variable-length non-contiguous regions (such as the exons of a gene).

### its_a_scam for single cell RNA and ATAC-seq count matrix generation and QC

`its_a_scam`, built on top of `gcat`, is a suite of tools for count matrix generation and per-cell QC. `its_a_scam` provides programs that work with other tools (such as BWA MEM [9], samtools [10], and STARsolo [11]) for count matrix generation, and thereby enables building modular single cell analysis pipelines. The major contribution of `its_a_scam` is a set of tools for generating per-cell quality control metrics computed from the alignments rather than from the count matrix. Being agnostic to the underlying single cell RNA-seq and ATAC-seq protocol, `its_a_scam` enables uniform count matrix generation and QC for pooled single cell analysis.

### scATAC-seq count matrix generation

Pipelines for aligning and calling peaks in bulk ATAC-seq samples are well established [12]. Using these pipelines for single cell ATAC-seq analysis requires additional steps of keeping track of the cell barcode for each read. We facilitate this process by providing programs for adding the cell barcode as a part of the fastq name (to be carried over to the aligned SAM file), barcode matching for 10x Multiome datasets, filtering low quality cells from a SAM file, generating count matrices, and generating a fragments file (Table 2: scATAC-seq count matrix generation). We evaluated the accuracy of `its_a_scam` using the ‘peripheral blood mononuclear cells (PBMC) from a Healthy Donor – Granulocytes Removed Through Cell Sorting (3k) Multiome’ dataset from 10x Genomics [13]. The fragments file generated using `its_a_scam` tools together with BWA [9] and samtools [10] is highly concordant with the one generated with CellRanger (Supplementary Fig 1). They identify nearly identical cells (Jaccard index: 0.996), and have high correlations for the counts in common peaks (see Methods) per cell (*R* ≈ 1), number of peaks per cell (*R* ≈ 1), and cell-peak counts (Median Spearman’s correlation: 0.99).

**Table 2.**
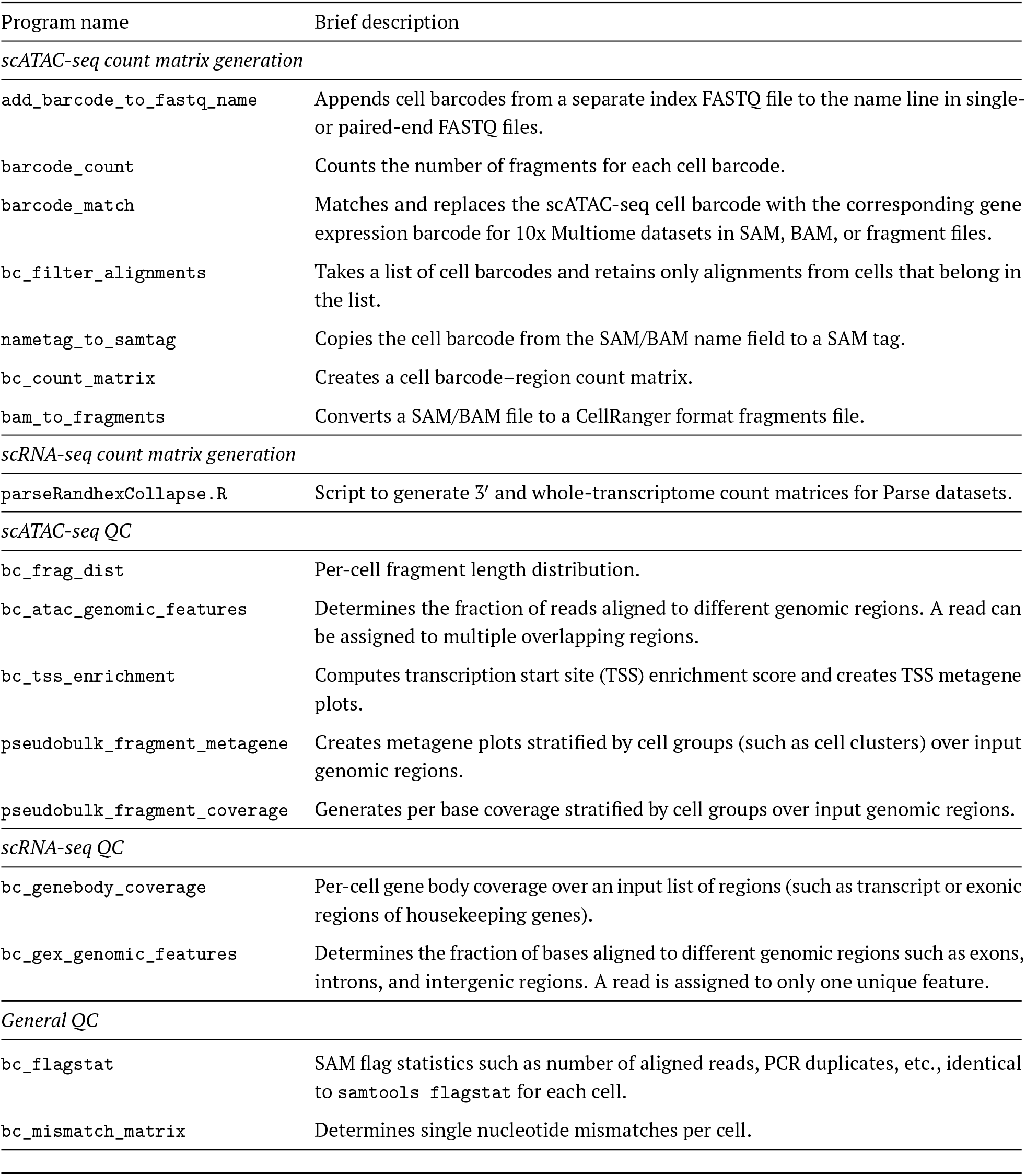
`its_a_scam` **is a collection of tools for pre-processing and QC of single cell data**.

**Figure 1.**
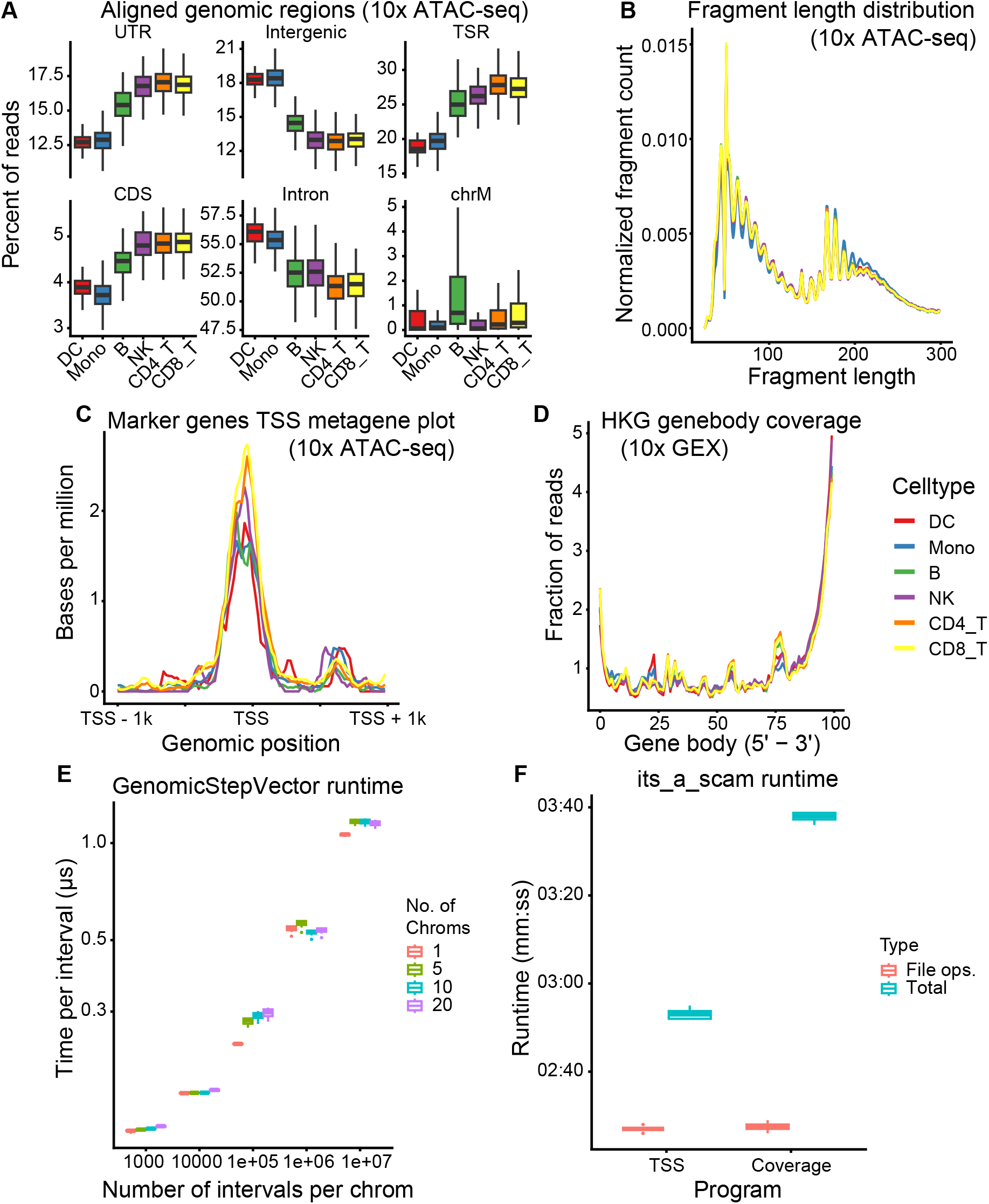
Quality control with and benchmarking of its_a_scam and gcat. **(A)** Aligned genomic regions of 10x scATAC-seq data. Transcription start regions (TSR) are 1000 bp upstream of the transcription start site (TSS). CDS: coding sequence; UTR: untranslated region; DC: dendritic cells; Mono: monocytes; NK: natural killer cells. **(B)** Fragment length distribution of 10x scATAC-seq data. **(C)** Metagene plot of nucleosome-free reads at the regions flanking the TSS (TSS ± 1kb) of T cell marker genes. **(D)** Genebody coverage over housekeeping genes (HKG) for 10x scRNA-seq data. **(E)** Time per insertion into a `GenomicStepVector<int=` stratified by the number of chromosomes. **(F)** Runtime of `its_a_scam` programs that make extensive use of `GenomicStepVector`.

### scRNA-seq count matrix generation

STARsolo [11] is a versatile open-source tool for gene expression quantification that is easily adaptable to a wide range of scRNA-seq protocols. It is designed to be used out of the box for 10x Genomics datasets, but its parameters vary depending on the 10x protocol used. We therefore do not duplicate efforts and simply provide a brief tutorial of the commonly used parameters for 10x 3^′^, 5^′^, and Multiome datasets. In contrast, STARsolo’s output for Parse Biosciences whole-transcriptome alignments requires post-processing. The Parse Biosciences protocol uses a combinatorial barcoding approach [14, 15] to uniquely tag cells without the need for extensive instrumentation. STARsolo’s output for these reads with “complex” barcodes requires random-hexamer collapsing to generate the final count matrix [16]. `its_a_scam` provides a program to generate both 3^′^ and whole-transcriptome count matrices from these STARsolo outputs (Table 2: scRNA-seq count matrix generation). We evaluated the accuracy of the whole-transcriptome counts generated with STARsolo and `its_a_scam` using the Parse H1 and H2 datasets from [17]. These count matrices are concordant with the ones generated using the Parse pipeline (split-pipe). They identify nearly identical cells (Jaccard index: ≥ 9.96), and have high correlations for the gene counts per cell (*R* ≥ 0.98) and number of genes per cell (*R* ≥ 0.97). For consistency, we counted genes analogously to other protocols (see Methods), leading to an overall slightly decreased median Spearman’s correlation of the cell-gene count vectors (≥ 0.88; Supplementary Figs 2 and 3).

#### QC of scRNA-seq and scATAC-seq data with **its_a_scam**

`its_a_scam` provides tools for generating QC metrics directly from aligned scRNA-seq and scATAC-seq BAM files (summarized in Table 2). These metrics complement the typical QC metrics generated from count matrices and can aid both in quality control and in comparative analysis post-clustering. `its_a_scam` typically reports the QC metric per cell barcode, which can be used downstream at multiple levels: (1) at the sample level to understand sample-specific properties, (2) at the cluster level to identify differences between clusters (or celltypes after annotation), or (3) at the single cell level for per-cell metrics. `its_a_scam` separates the code to generate the QC metrics and the code to visualize them. The programs typically take the SAM/BAM file and the list of barcodes as input to generate a TSV file with the per-barcode QC metrics. We provide separate R scripts to visualize the QC metrics across all levels. In addition, a user can easily adapt or create scripts for custom visualization from the TSV files.

We demonstrate the use of `its_a_scam` on the 10x PBMC Multiome dataset. At the sample level, it contains a high fraction of reads aligned to the transcription start region (transcription start site [TSS] − 1kb; TSR) and a low fraction to the mitochondrial chromosome (Supplementary Fig 5a). It exhibits the expected periodic fragment length distribution with peaks corresponding to nucleosome-free and nucleosome-bound fragments (Supplementary Fig 6a). The cells were then jointly clustered and annotated into six celltypes (Supplementary Fig 4). Per-cell and cluster QC metrics show that NK and T cells have fewer scATAC-seq reads per cell but similar mapping and duplication rates compared with the other celltypes (Supplementary Fig 9). The monocytes and dendritic cells have a lower fraction of their reads aligned to TSR, untranslated region (UTR), and coding sequence (CDS) regions, and a higher fraction to the intronic and intergenic regions, while the B, NK, and T celltypes exhibit the opposite trend (Figure 1a). All celltypes exhibit an almost identical fragment length distribution and TSS enrichment score (Figure 1b and Supplementary Fig 6b, Supplementary Fig 7). The metagene plot of nucleosome-free reads (≤ 150bp) at the regions flanking the TSS (TSS ± 1kb) of T cell marker genes shows higher accessibility for the T cells clusters, while all celltypes have similar accessibility around the TSS of house-keeping genes (Figure 1c and Supplementary Fig 8). The gene expression reads show similar genebody coverage over all house-keeping genes across celltypes (Figure 1d and Supplementary Fig 11), while monocytes have a lower fraction of bases aligned to the CDS and UTR regions and a higher fraction to the intronic regions compared with the other celltypes (Supplementary Fig 10). These metrics, when viewed along with the celltype annotation, reveal celltype-specific differences that cannot be obtained from the count matrices alone.

### Benchmarking GenomicStepVector, the workhorse of gcat and its_a_scam

We benchmarked the efficiency of storing data in and retrieving data from`GenomicStepVector` using nanobench [18]. `its_a_scam` can take aligned SAM/BAM files that are either unsorted or sorted by genomic coordinates (for example, using `samtools sort`). To mimic these conditions, we inserted 10^3^–10^7^ intervals (span of 10^7^ bases with a mean length of 500 bases) into a `GenomicStepVector`. At 10^3^ intervals, corresponding to *∼* 0.05× coverage, insertion of sorted intervals is 1.28× faster than unsorted intervals, and at 10^7^ intervals (*∼* 500× coverage) it is 28.6× faster (Supplementary Fig 12a). Furthermore, insertion time per interval is essentially independent of the number of chromosomes (Figure 1e). After all intervals are inserted, the time per query is not impacted by the number of queries, and the query intervals sorted by genomic positions are *∼* 3× faster than unsorted queries (Supplementary Fig 12b). The time per query increases as the number of intervals in the data structure increases, reflecting the larger number of steps spanning the queried region (Supplementary Fig 12c). Aligned BAM files are typically coordinate sorted, which makes `its_a_scam` programs faster. We evaluated the performance of `GenomicStepVector` on real data (10x Multiome PBMC dataset) by benchmarking the `its_a_scam` programs that use it. `bc_tss_enrichment` uses a `GenomicStepVector` to keep track of the TSS regions, predominantly querying the data structure. The total runtime for a dataset with 79.9M alignments was 173.6 ± 1.1s with 147.5 ± 0.6s (84.9%) consumed for file operations (Figure 1f TSS). `pseudobulk_fragment_coverage` uses two `GenomicStepVectors`, one to query whether an alignment belongs to a desired region and another to keep track of the coverage, thereby exercising both querying and insertion into the data structure. The total runtime for the same dataset was 218.5 ± 1.0s with 148.0 ± 0.9s (67.7%) for file operations (Figure 1f Coverage). The peak memory consumed by these programs is small, 236.5MiB and 3.12GiB, respectively (Supplementary Fig 12d). Taken together, `GenomicStepVector` is an efficient data structure for storing data associated with genomic coordinates, and is fast enough not to be the bottleneck for programs in `its_a_scam`.

## Discussion

We have introduced `its_a_scam`, a suite of tools for modular analysis of single cell RNA-seq and ATAC-seq data. We leveraged `its_a_scam` together with BWA MEM and samtools to show that the output is concordant with that of CellRanger. The modular nature of `its_a_scam` allows a user to easily adopt it into their own pipelines, for example, by using other tools such as Bowtie2 [19], Picard [20], or HMMRATAC [21]. Furthermore, the Parse Biosciences whole-transcriptome count matrices generated using `its_a_scam` together with STARsolo are concordant with those of split-pipe. The single cell RNA-seq and ATAC-seq QC metrics generated directly from the alignment files enable analysis beyond just using the metrics for filtering low quality cells, enabling comparative analysis of alignment differences in celltypes after annotation. `its_a_scam` can also be used for QC of data that has been generated by other upstream pipelines such as CellRanger or chromap [22], as long as they output data in SAM format. Taken together, `its_a_scam` enables modular single cell count matrix generation, quality control, and analysis using open source tools.

We also introduce `gcat`, a C++ library for parsing genomic data files and working with genomic coordinates. Here, we used `gcat` to build `its_a_scam`, but it is by no means limited to `its_a_scam`. The `gcat` library can be used to develop other analysis or quality control programs as required by the user. Specifically, the `GenomicStepVector` combined with genomic file parsing makes `gcat` a powerful library. For example, `gcat` can be used for developing programs for coverage or variant analysis, and gene expression quantification.

The key limitations at the moment are: (1) Neither `gcat` nor `its_a_scam` offers support for multi-threading. However, as shown by our benchmarking, single-threaded `its_a_scam` is fast and would not bottleneck the analysis. (2) `its_a_scam` currently requires SAM/BAM files as input and cannot directly work with fragments files. (3) `its_a_scam` does not offer programs for analyzing spatial genomic data. These will be addressed in subsequent updates to the tools.

Overall, `gcat` and `its_a_scam` provide tools for building both custom analysis programs and modular pipelines for genomic data analysis. The documentation and usage instructions for `gcat` and `its_a_scam` are available at https://gcatlib.readthedocs.io/ and https://its-a-scam.readthedocs.io/, respectively. We provide a detailed description for a user to generate count matrices and QC single cell RNA-seq and ATAC-seq datasets starting from FASTQ files. Of course, they are open source (licensed under GNU GPL version 2), enabling users to adapt and build on our tools as desired.

## Material and Methods

### Datasets

#### 10x Multiome

PBMC from a Healthy Donor - Granulocytes Removed Through Cell Sorting (3k) analyzed using Cell-Ranger ARC 2.0.0 (Date Published: 2021-05-03).

#### Parse Biosciences

The parse_H1 and parse_H2 datasets from [17] (GEO accession number GSE246624).

#### Gene lists

We used the list of human house-keeping (HK) genes as defined by Eisenberg & Levanon [23]; we mapped the gene identifiers from RefSeq to Ensembl gene identifiers [24], yielding 4,211 unique genes.

### Read alignment and count matrix generation

#### Aligning 10x ATAC-seq reads

The cell barcodes from the barcode fastq file were added to the name field of the paired-end ATAC fragment fastq files using `add_barcode_to_fastq_name`. The paired-end reads were aligned to the human reference genome (GRCh38 primary assembly) using `BWA MEM` (version: 0.7.17-r1188). Supplementary alignments and PCR duplicates were removed using `samtools` (version: 1.17). The ATAC-seq cell barcode was matched and replaced with the respective GEX cell barcode using `barcode_match`. The number of fragments per cell was counted using `barcode_count`. Cells with fewer than 1000 fragments were discarded with `bc_filter_alignments`. In addition, the cell barcode matched BAM file was converted to fragments file using `bam_to_fragments`. The detailed procedure for using `its_a_scam` for scATAC-seq analysis is described here https://its-a-scam.readthedocs.io/en/latest/10x_atac_seq/.

#### Aligning 10x RNA-seq reads

The 10x scRNA-seq reads were aligned to the human reference genome (GRCh38 primary assembly) with Gencode gene annotation (v44) using STARsolo (version: 2.7.11a) with the following parameters:

~~~
--readFilesCommand zcat --soloType CB_UMI_Simple --soloUMIlen 12
--outSAMattributes NH HI nM AS CR UR CB UB GX GN sS sQ sM
--outSAMtype BAM SortedByCoordinate --outFilterScoreMin 30
--soloCBwhitelist 737K-arc-v1.txt
--clipAdapterType CellRanger4 --soloCBmatchWLtype 1MM_multi_Nbase_pseudocounts
--soloUMIfiltering MultiGeneUMI_CR --soloUMIdedup 1MM_CR
--soloCellFilter EmptyDrops_CR --soloMultiMappers EM --soloBarcodeReadLength 1
--soloFeatures GeneFull GeneFull_ExonOverIntron GeneFull_Ex50pAS Gene Velocyto
~~~

The parameter choices for aligning 10x datasets with STARsolo are described here https://its-a-scam.readthedocs.io/en/latest/10x_gex/

#### Aligning Parse Biosciences RNA-seq reads

Each sub-library of the Parse Biosciences WT was aligned to the human reference genome (GRCh38 primary assembly) with Gencode gene annotation (v44) using STARsolo with the following parameters:

~~~
--readFilesCommand zcat --outSAMtype BAM SortedByCoordinate
--outSAMattributes NH HI nM AS CR UR CB UB GX GN sS sQ sM
--soloCellFilter EmptyDrops_CR --outFilterScoreMin 30
--soloCBmatchWLtype EditDist_2 --soloUMIfiltering MultiGeneUMI_CR
--soloUMIdedup 1MM_CR --soloFeatures GeneFull GeneFull_ExonOverIntron GeneFull_Ex50pAS
--soloMultiMappers EM --soloBarcodeReadLength 0 --soloType CB_UMI_Complex
--soloCBwhitelist bc_data_v1.txt bc_data_v1.txt bc_data_n99_v5.txt--soloCBposition 0_10_0_17 0_48_0_55 0_78_0_85 --soloUMIposition 0_0_0_9
--clipAdapterType CellRanger4 --clip5pAdapterSeq AAGCAGTGGTATCAACGCAGAGTGAATGGG
~~~

Random hexamers were collapsed and Seurat objects were generated using `parseRandhexCollapse.R`. The sub-libraries were merged into one SeuratObject using the Seurat `merge` function (version: 5.0.3) [25]. The procedure for using `its_a_scam` to generate the 3^′^ and random-hexamer collapsed count matrices is described here https://its-a-scam.readthedocs.io/en/latest/parse_wt/

### Evaluating the concordance of single cell analyses using its_a_scam

#### Concordance of its_a_scam pipeline for 10x ATAC-seq data

The FASTQ files from the 10x Multiome PBMC from a Healthy Donor - Granulocytes Removed Through Cell Sorting (3k) dataset were used to generate the fragments file using `its_a_scam` as described above. Peaks were called using `CallPeaks` in Signac [26], which internally calls MACS3 (version: 3.0.1) [27]. The CellRanger fragments file for comparison was downloaded directly from the 10x datasets page. To keep the comparison consistent, the peaks called with the `its_a_scam` fragments file were used to generate both the `its_a_scam` and CellRanger count matrices.

#### Concordance of its_a_scam pipeline for Parse Biosciences RNA-seq data

The FASTQ files were downloaded from GEO (GSE246624) and processed as described above to generate the count matrix using STARsolo and `its_a_scam`. The count matrices generated using Parse split-pipe were downloaded from GEO, and loaded using Seurat for comparison.

### 10x multiome data cell clustering, annotation, and QC

#### Cell clustering

The 10x Multiome gene expression (GEX) and ATAC-seq reads were processed using `its_a_scam` as described above. The GEX count matrix generated with the `GeneFull_Ex50pAS` model was used. Cells that had both GEX and ATAC-seq barcodes were used for downstream analysis. Peaks were called using MACS3 (version: 3.0.1) and the joint count matrices were generated with Seurat. The weighted nearest neighbor graph was constructed using `FindMultiModalNeighbors`, and clusters were generated using `FindClusters` with the smart local moving (SLM) algorithm on the wsnn graph at a resolution of 0.4. The joint clustering generated 10 clusters.

#### PBMC celltype annotation

The PBMC celltype marker genes were taken from the HuBMAP Azimuth database (Human PBMC celltype.l1) [28]. The module scores were generated for each celltype using `AddModuleScore`. The module scores clearly separated 8 of the 10 clusters, and these were annotated as the celltype with the highest score. Cluster 3 and 5 had high module scores for multiple celltypes. We examined the key PBMC marker genes and annotated these clusters as CD8 T cells given the high expression of CD8A and CD8B (Supplementary Fig 4). Multiple clusters annotated as the same celltype were merged for downstream analysis.

#### QC metric generation and visualization

Alignments with a mapping quality higher than 30 were used to generate the QC metrics. The metrics were visualized using the plotting scripts supplied with `its_a_scam`. All QC metrics were generated using the default parameters. The metagene plots were generated with `pseudobulk_fragment_metagene` for nucleosome-free fragments (fragment length ≤ 150bp). T cell marker genes for the metagene plot were taken as the union of CD4 and CD8 T cell marker genes.

### Benchmarking **gcat** **and** **its_a_scam**

#### Benchmarking hardware

All benchmarking was performed on a workstation with an AMD Ryzen 9 5900X processor containing 96GB memory. The system was set to performance mode using `pyperf system tune` prior to benchmarking. The benchmark program was pinned to one processor using taskset. The tests were performed after a reboot with only necessary user programs running.

#### Inserting into **GenomicStepVector**

A `GenomicStepVector<int=` was used. The starting position of the interval to be inserted was sampled from a uniform distribution (*U*[0, 10^7^]), and the length of the intervals are sampled from *U*[1, 10^3^]. The required number of intervals were pre-generated with a constant insertion value of 1. If needed, the intervals were sorted by genomic positions prior to insertion. The insertions were then benchmarked 10 times after 3 warm-up runs.

#### Querying **GenomicStepVector**

The starting position of the genomic interval was sampled from a uniform distribution (*U*[0, 10^7^]), and the length of the interval was sampled from *U*[1, 10^3^]. The required number of intervals were generated and inserted into a `GenomicStepVector<int=` with a fixed value of 1. The required number of query intervals were then generated from the same distribution. If needed, the query intervals were sorted by genomic positions prior to querying. The queries were then benchmarked 10 times after 3 warm-up runs.

#### Benchmarking **its_a_scam**

The `bc_tss_enrichment` and `pseudobulk_fragment_coverage` from `its_a_scam` were benchmarked using the 10x Multiome PBMC from a Healthy Donor dataset. The BAM file was pre-filtered using `bc_filter_alignments` to retain alignments that belong to the valid cells and have a mapping quality of at least 30. 79,906,518 alignments from 2,628 cells were retained. Each program was run 10 times with the default parameters, and the wall-clock time and peak memory usage were determined using `/usr/bin/time -v`. The “File ops.” measurements were taken using an invalid cell barcode and a 1 base region, and thus minimizing time spent on data processing and maximizing the time spent on file reading and parsing.

## Author contributions

Conceptualization: RKP, HVM; Methodology: RKP; Software: RKP; Validation: RKP; Formal analysis: RKP; Investigation: RKP; Resources: RKP; Writing - original draft: RKP; Writing - review & editing: RKP, HVM; Visualization: RKP; Supervision: RKP, HVM; Project administration: HVM; Funding acquisition: HVM.

## Data availability

The ‘PBMC from a Healthy Donor - Granulocytes Removed Through Cell Sorting (3k)’ analyzed using CellRanger ARC 2.0.0 (Date Published: 2021-05-03) was downloaded from the 10x datasets page. The parse datasets from [17] were downloaded from GEO (GSE246624).

## Code availability

`gcat` (distributed as gcatlib) is available at https://github.com/rishvanth-kp/gcatlib. The benchmarking code is located in the `tests` directory. The installation instructions and class interface documentation are available at https://gcatlib.readthedocs.io/.

`its_a_scam` is available at https://github.com/rishvanth-kp/its_a_scam. The plotting scripts are in the `scripts` directory. The installation instructions, program usage instructions, and documentation for count matrix generation and QC are available at https://its-a-scam.readthedocs.io/.

Custom analysis code for joint clustering and annotation, parsing and plotting the benchmarking results, and concor-dance analysis were written in R (version ≥ 4.1.2). All analyses code is freely available on GitHub: https://github.com/meyer-lab-cshl/gcat_its_a_scam_analysis.

