## Supplementary Figures for "gcat, its_a_scam: modular tools for single cell data processing and per-cell quality control"

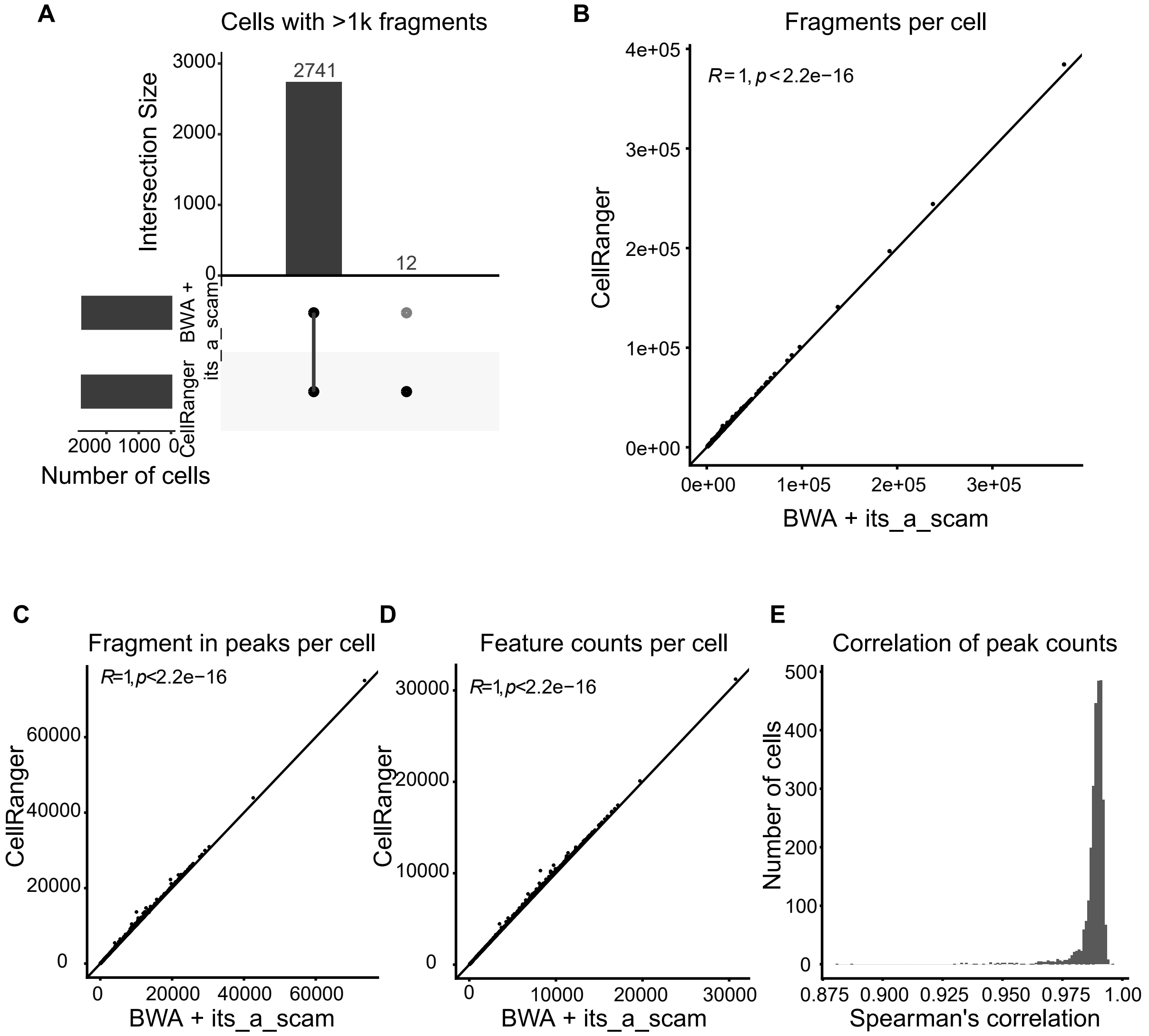


**Supplementary Figure 1. 10x ATAC-seq `its_a_scam` pipeline is concordant with CellRanger. (A)** Number of common cells identified by the two pipelines. **(B)** Fragments per cell (Pearson correlation is indicated). **(C)** Fragments in peaks per cell (Pearson correlation is indicated). **(D)** Features (peak) counts per cell (Pearson correlation is indicated). **(E)** Spearman's correlation of per-cell feature (peak count) vectors.

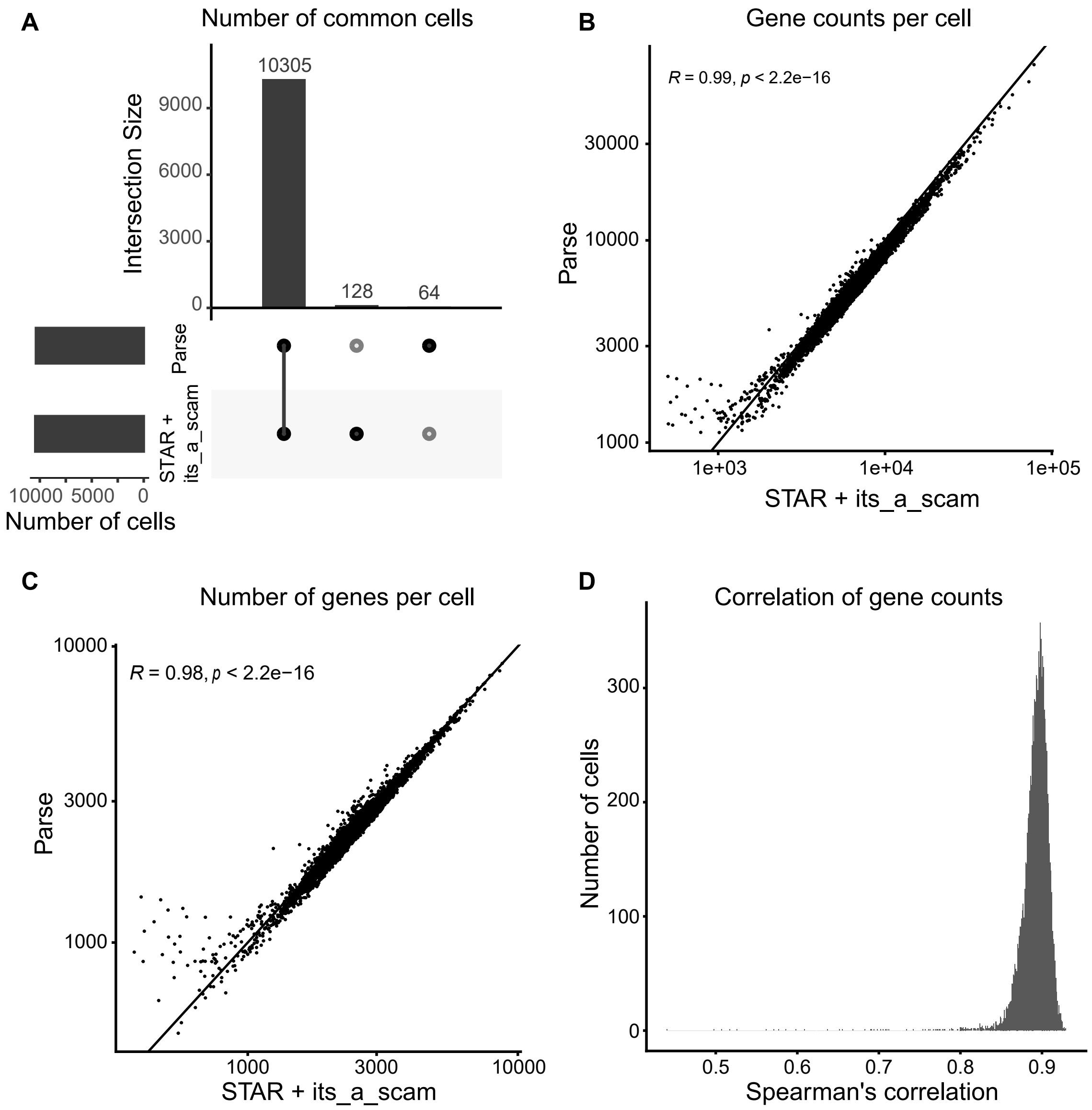


**Supplementary Figure 2. Parse Bioscience WT `its_a_scam` pipeline is concordant with the Parse pipeline split-pipe (H1 dataset). (A)** Number of common cells identified by the two pipelines. **(B)** Gene counts per cell (Pearson correlation is indicated). **(C)** Number of genes per cell (Pearson correlation is indicated). **(D)** Spearman's correlation of per cell gene counts.

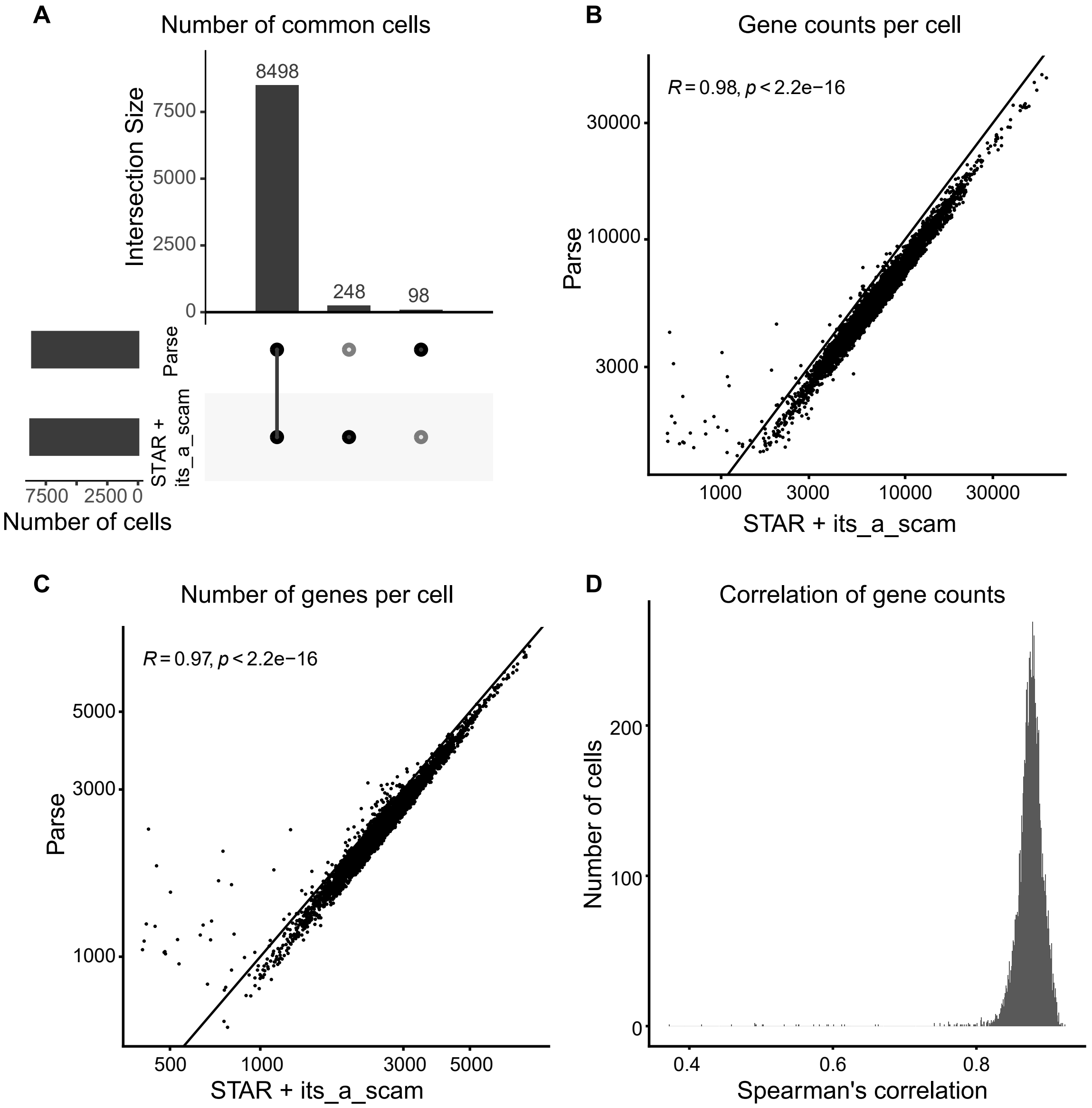


**Supplementary Figure 3. Parse Bioscience WT `its_a_scam` pipeline is concordant with the Parse pipeline split-pipe (H2 dataset). (A)** Number of common cells identified by the two pipelines. **(B)** Gene counts per cell (Pearson correlation is indicated). **(C)** Number of genes per cell (Pearson correlation is indicated). **(D)** Spearman's correlation of per cell gene counts.

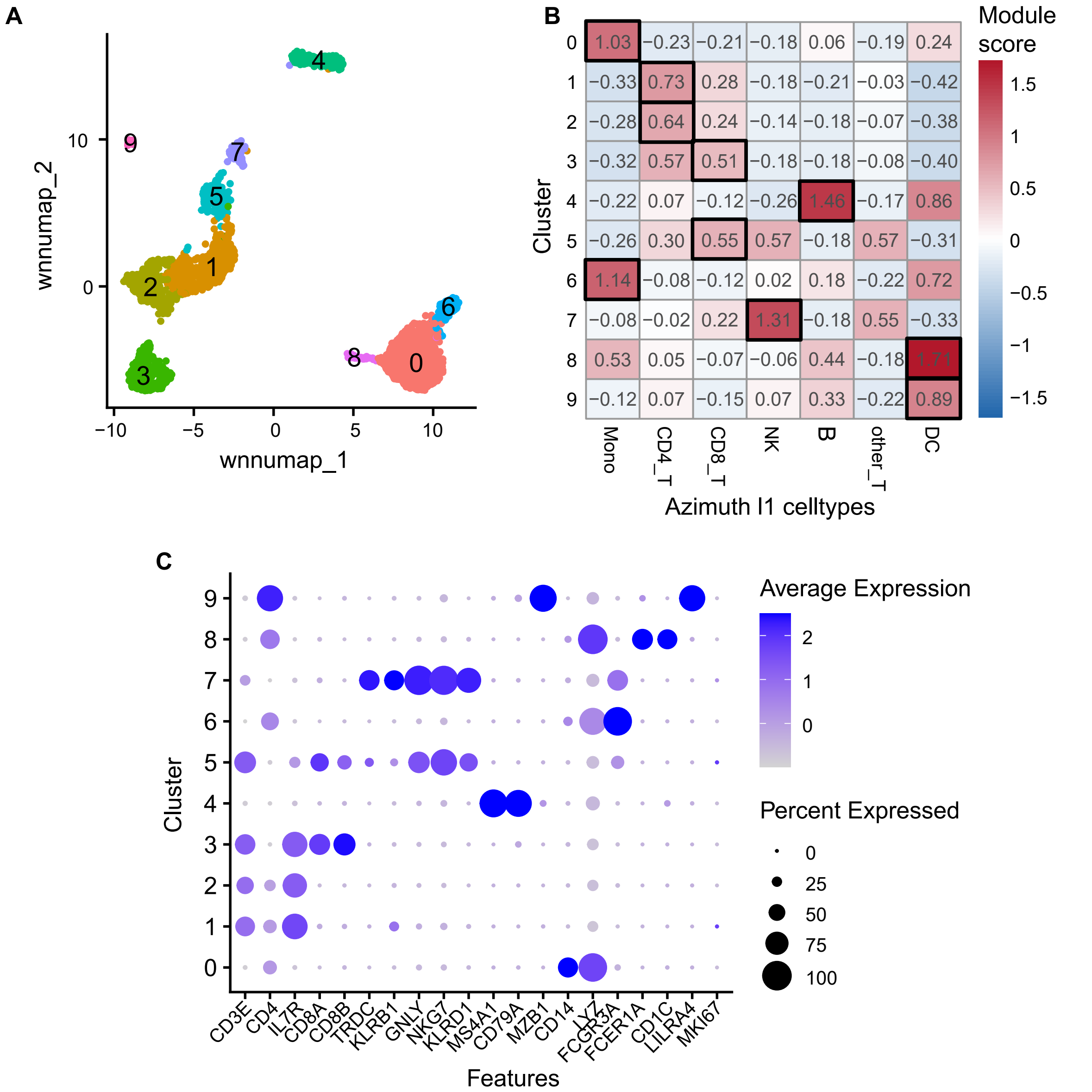


**Supplementary Figure 4. Joint clustering and annotation of 10x PBMC Multiome dataset. (A)** UMAP of single-nucleus RNA-seq and ATAC-seq jointly clustered cells with the cluster IDs labeled. **(B)** Module scores for the PBMC celltype marker genes were taken from the HuBMAP Azimuth database (Human PBMC celltype.l1). The bold box indicates the celltype annotation for each cluster. **(C)** Expression of key PBMC marker genes by cluster. We annotated clusters 3 and 5 as CD8_T cells given the high expression of CD8A and CD8B.

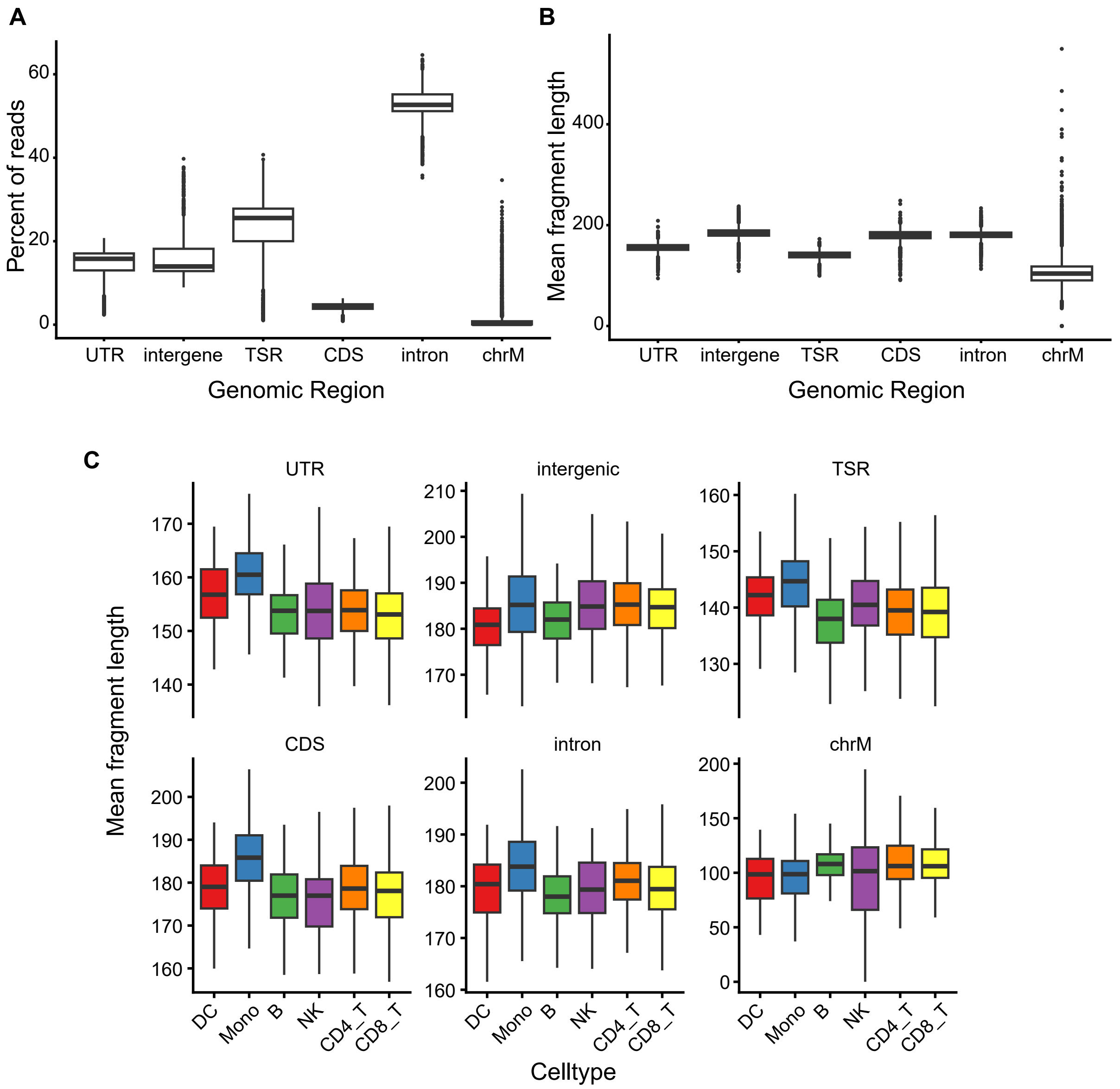


**Supplementary Figure 5. 10x ATAC-seq aligned genomic regions generated with `bc_atac_genomic_features`. (A)** Sample level fraction of aligned reads in different genomic regions. **(B)** Sample level mean fragment length in different genomic regions. **(C)** Celltype level mean fragment length in different genomic regions. TSR: transcription start region; CDS: coding sequence; UTR: untranslated region; DC: dendritic cells; Mono: monocytes; NK: natural killer cells.

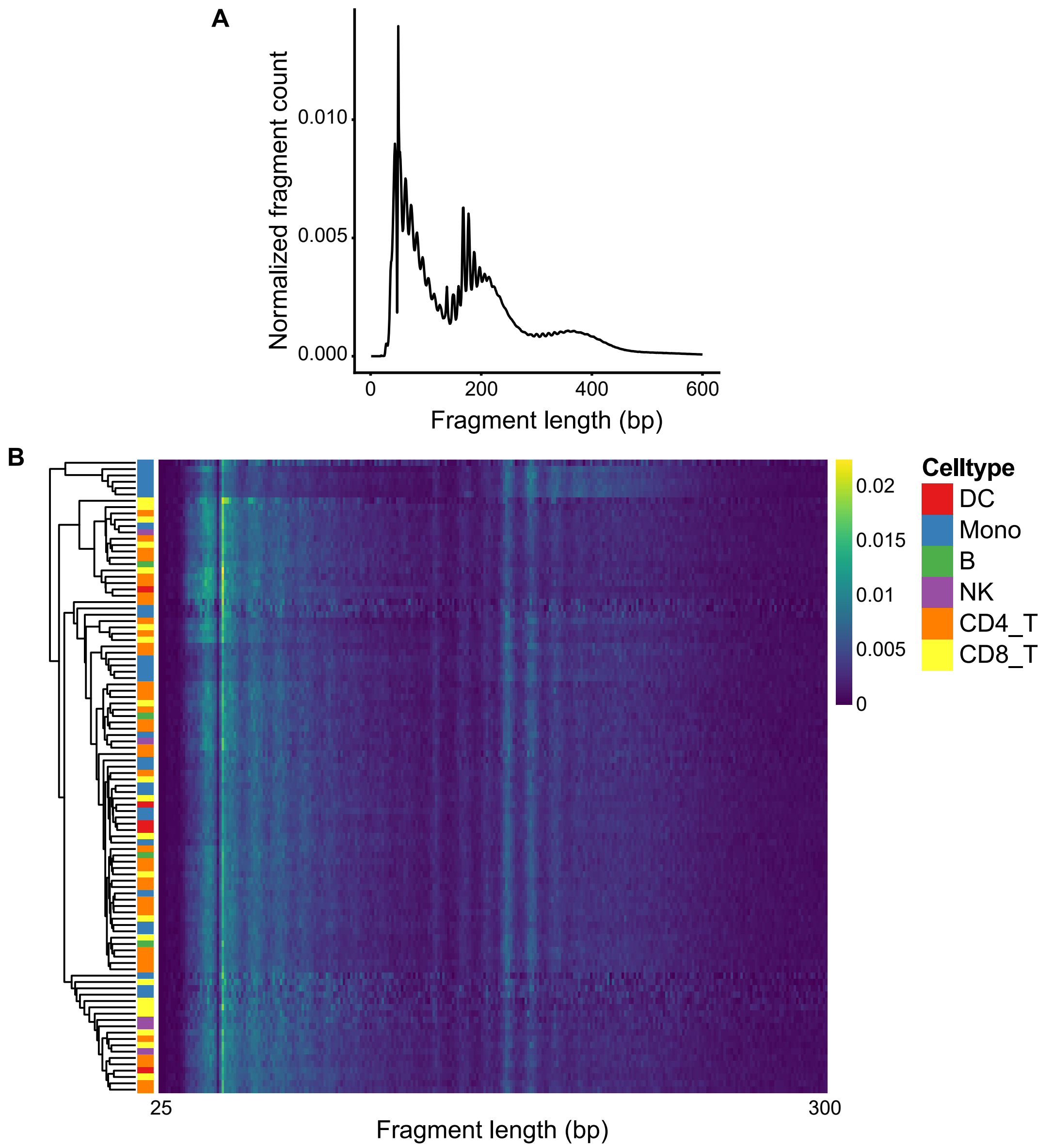


**Supplementary Figure 6. 10x ATAC-seq fragment length distribution generated with `bc_frag_dist`. (A)** Sample level fragment length distribution. **(B)** Per cell fragment length distribution. Heatmap is downsampled to 100 cells for clarity. DC: dendritic cells; Mono: monocytes; NK: natural killer cells.

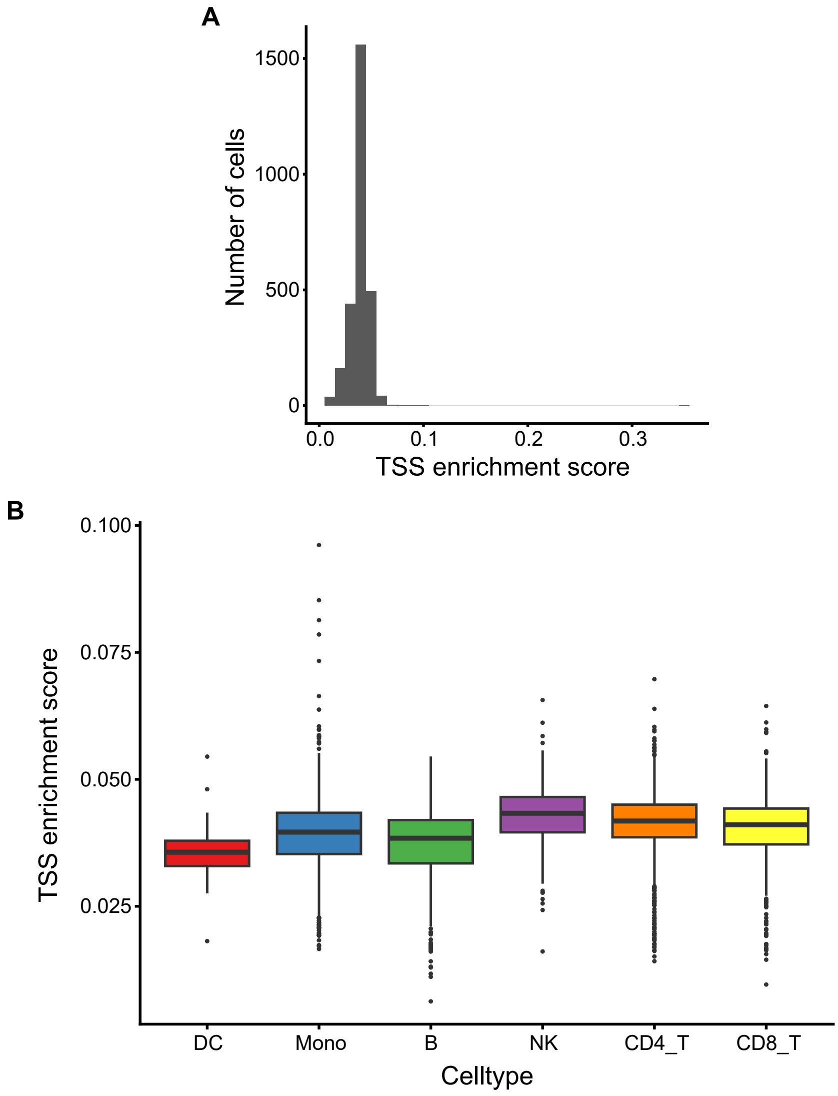


**Supplementary Figure 7. 10x ATAC-seq transcription start site (TSS) enrichment score generated with `bc_tss_enrichment`.** **(A)** Sample level TSS enrichment score. **(B)** TSS enrichment score stratified by celltype. DC: dendritic cells; Mono: monocytes; NK: natural killer cells.

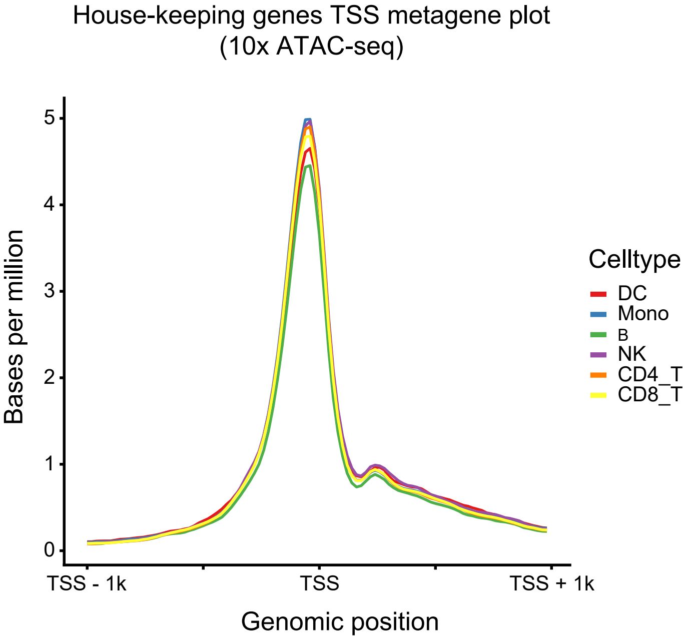


**Supplementary Figure 8. 10x ATAC-seq metagene plot generated with `pseudobulk_fragment_metagene`.** Metagene plot of nucleosome-free reads ($\leq 150$bp) at the regions flanking the transcription start site (TSS; TSS $\pm$ 1kb) of house-keeping genes. DC: dendritic cells; Mono: monocytes; NK: natural killer cells.

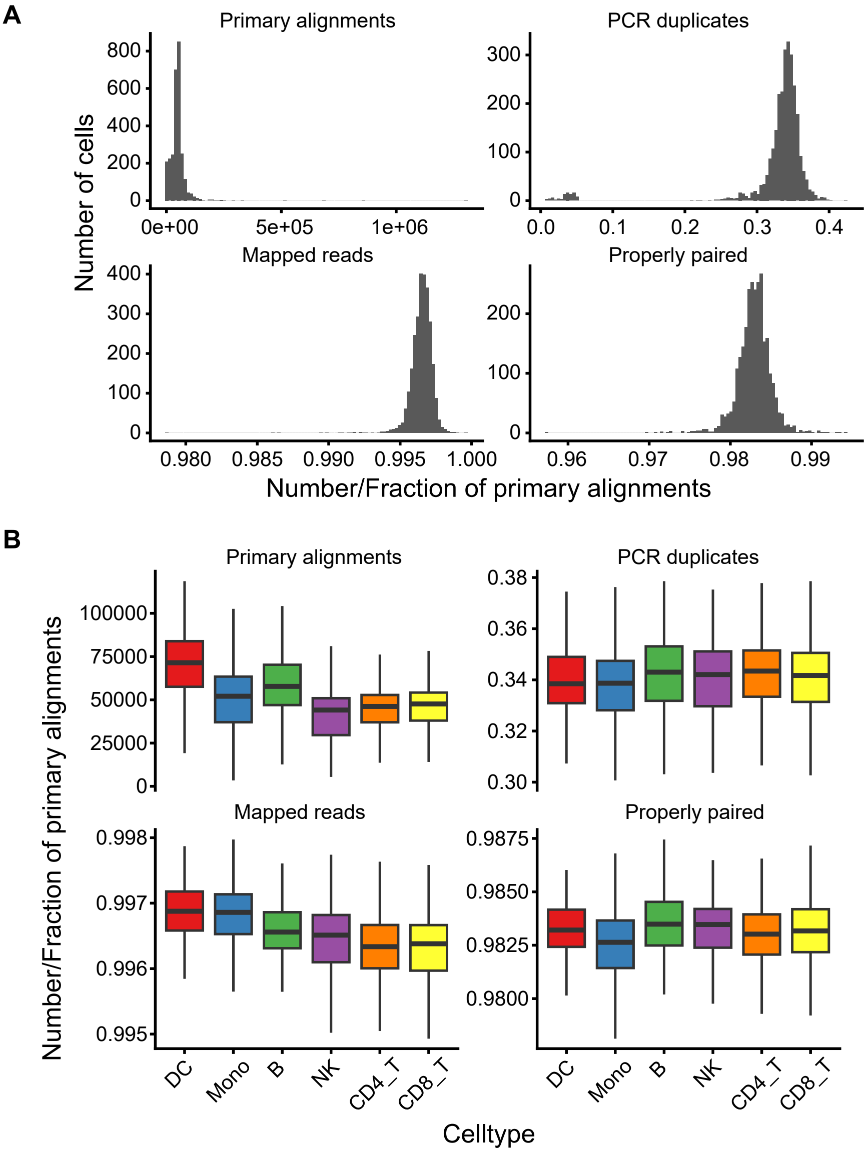


**Supplementary Figure 9. 10x ATAC-seq alignment statistics generated with `bc_flagstat`. (A)** Sample level alignment statistics. **(B)** Alignment statistics stratified by celltype. DC: dendritic cells; Mono: monocytes; NK: natural killer cells.

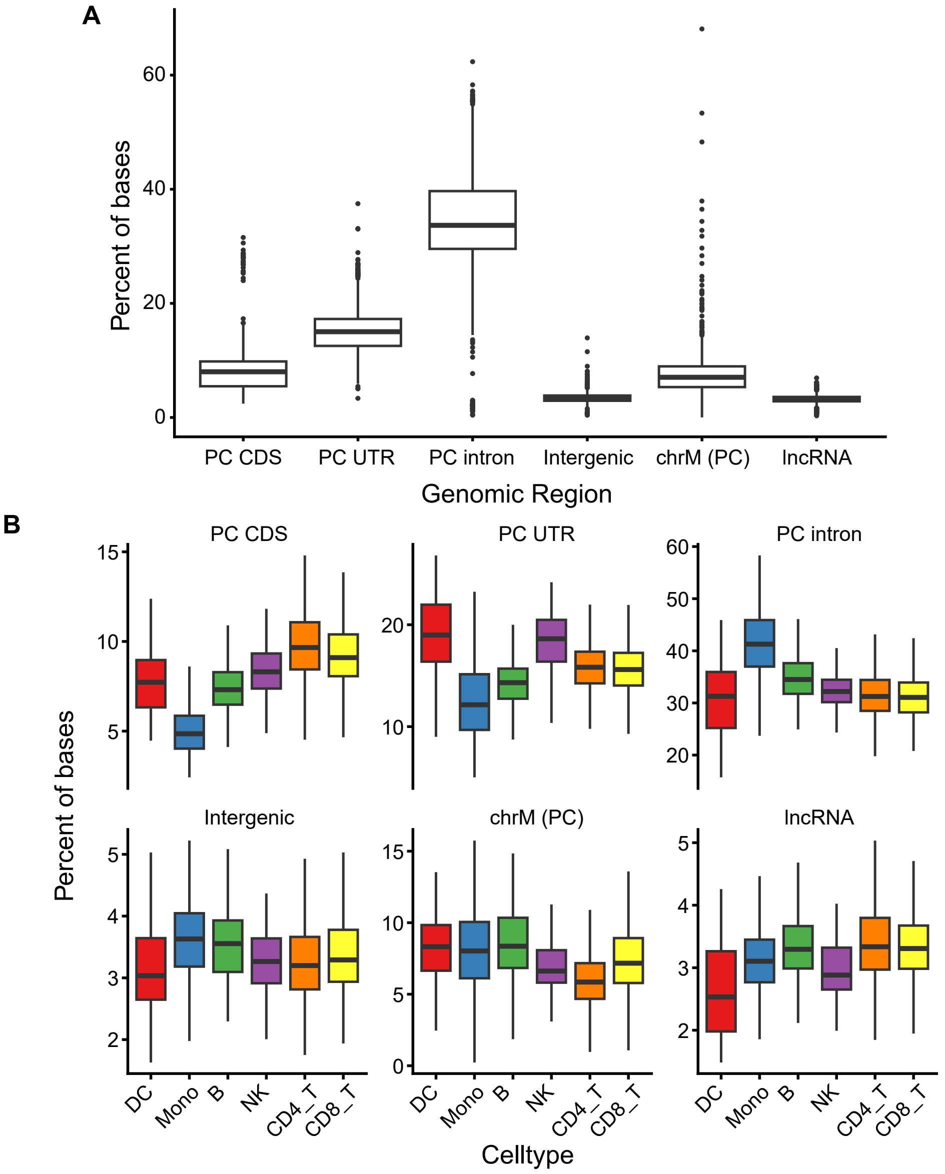


**Supplementary Figure 10. 10x GEX aligned genomic regions generated with `bc_gex_genomic_features`. (A)** Sample level fraction of aligned reads in distinct genomic regions. **(B)** Aligned regions stratified by celltype. PC: protein coding. CDS: Coding sequence. UTR: Untranslated regions; lncRNA: long non-coding RNA; DC: dendritic cells; Mono: monocytes; NK: natural killer cells.

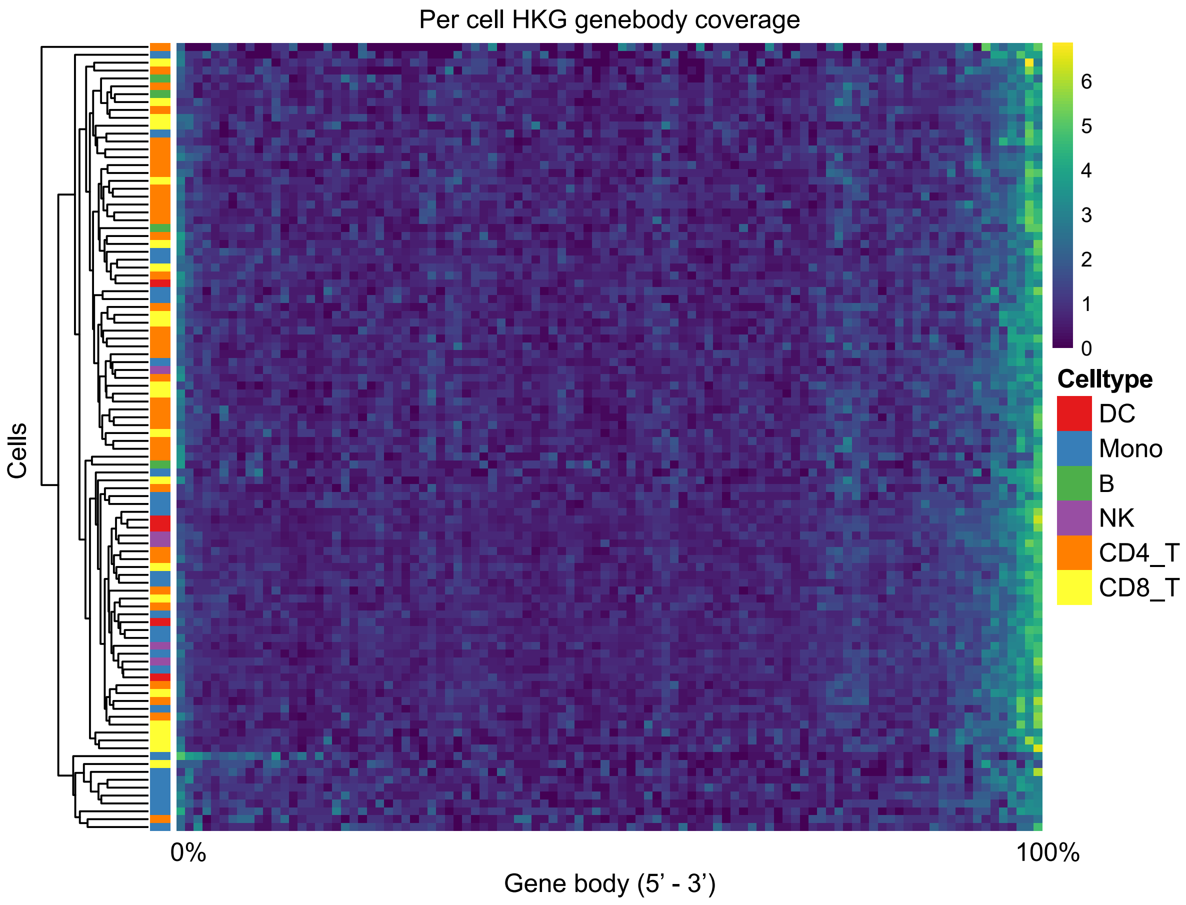


**Supplementary Figure 11. 10x GEX genebody coverage generated with `bc_genebody_coverage`.** Per cell genebody coverage over house-keeping gene (HKG) exons. Heatmap is downsampled to 100 cells for clarity. DC: dendritic cells; Mono: monocytes; NK: natural killer cells.

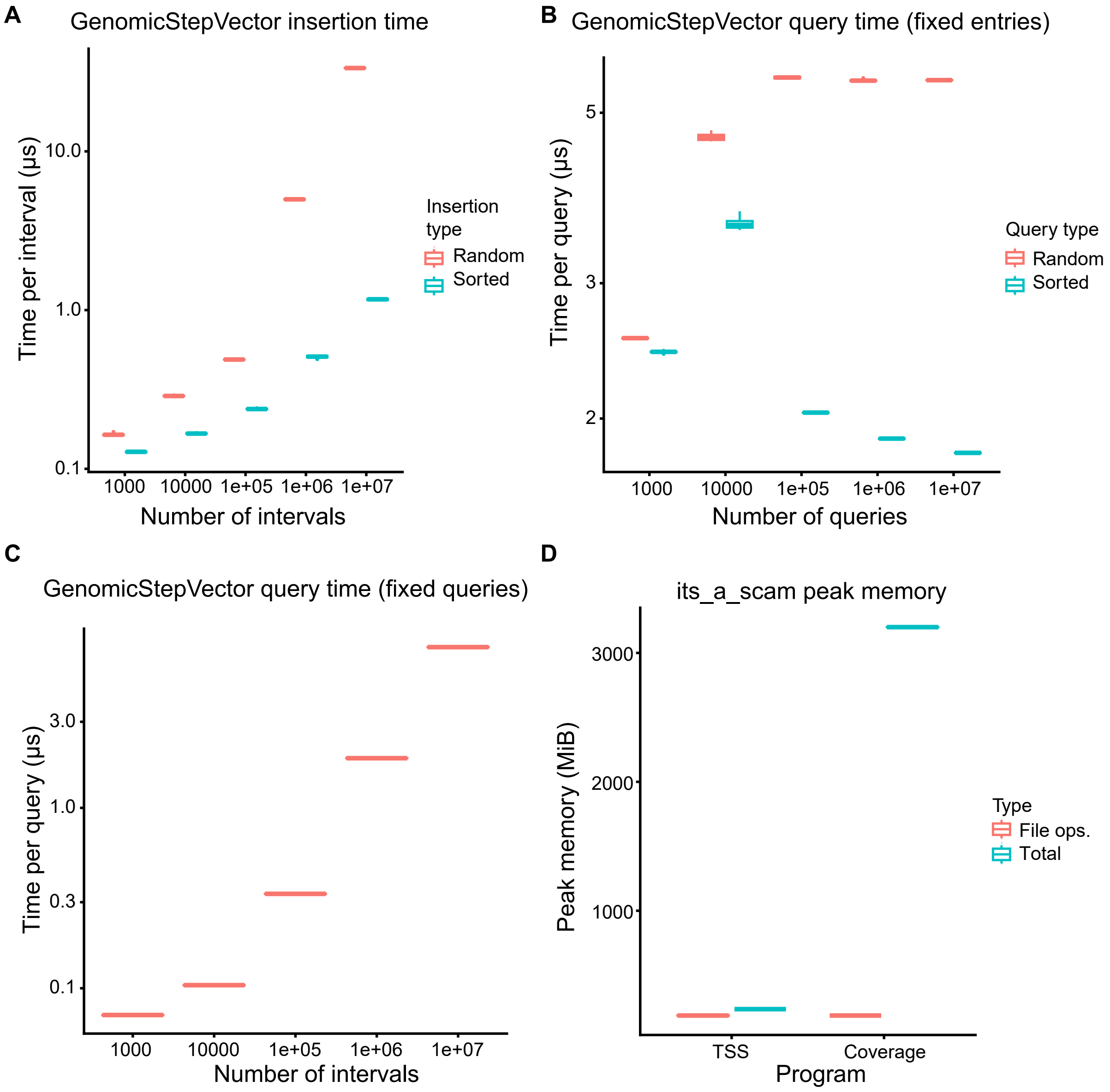


**Supplementary Figure 12. Benchmarking `gcat` and `its_a_scam`.** **(A)** Time per insertion into a single-chromosome `GenomicStepVector<int>` when the insertions are random or have sorted genomic coordinates. **(B)** Time per query into a single-chromosome `GenomicStepVector<int>` when the number of entries is fixed at 1M, and the number of queries is varied. **(C)** Time per query into a single-chromosome `GenomicStepVector<int>` when the number of queries is fixed at 1M, and the number of entries is varied. **(D)** Peak memory usage of `its_a_scam` programs that make extensive use of `GenomicStepVector`.
